# Making the Impossible Possible: Uniting TMT and DIA for Scalable Low-Input-Proteomics

**DOI:** 10.64898/2026.10.20.760427

**Authors:** Tim Thierer, Micha Johannes Birklbauer, Viktoria Dorfer, Karl Mechtler, Manuel Matzinger

**Affiliations:** Proteomics Tech Hub, Research Institute of Molecular Pathology (IMP), Vienna, Austria; Bioinformatics Research Group, University of Applied Sciences Upper Austria, Hagenberg, Austria; Gregor Mendel Institute of Molecular Plant Biology (GMI), Austrian Academy of Sciences, Vienna BioCenter (VBC), Vienna, Austria; Institute of Molecular Biotechnology (IMBA), Austrian Academy of Sciences, Vienna BioCenter (VBC), Vienna, Austria

**Keywords:** TMT, DIA, DDA, single-cell proteomics, mass spectrometry, liquid chromatography, Astral mass spectrometer

## Abstract

Isobaric multiplexing using tandem mass tags (TMT) and data-independent acquisition (DIA) are two powerful techniques in quantitative proteomics, each offering distinct advantages. Combining these strategies provides synergistic benefits for low-input applications such as single-cell proteomics (SCP). However, integrating these strategies has remained challenging due to co-isolation interference and increased data complexity. Here, we present a systematic, identification-linked evaluation of a hybrid DIA-TMTpro 18-plex workflow on the Orbitrap Astral mass spectrometer (MS). Using a two-proteome interference model of human and yeast digests, we benchmarked quantitative accuracy and precision of TMT based DDA and DIA acquisition strategies across a range of isolation windows, injection times and input amounts. We provide READ (**R**eporter Ion **E**xtractor and **A**nnotation **D**irector), an in-house developed Python-based tool which enables streamlined reporter ion (RI) quantification and co-isolation purity assessment. Using READ, TMT-based quantification becomes possible from DIA-focused search engines like Spectronaut or DIA-NN, which do not natively support TMT. We show that co-isolation interference drives both ratio compression and ratio inflation in TMT RI-based quantification regardless of acquisition mode. As an interference-resistant alternative, we further evaluated quantification via TMT complementary ions, which achieved near-label free quantitative accuracy at the cost of reduced plexing capacity and proteome depth. Together, these results establish practical guidelines for combining DIA and TMTpro on the Orbitrap Astral mass spectrometer and provide a foundation for extending DIA-TMT workflows to low-input applications such as SCP, where sensitivity, accuracy and throughput are critical.

## Introduction

Mass spectrometry (MS)-based proteomics has become an indispensable tool for understanding biological systems, now even at the single-cell level. The ability to study the proteome of single cells will lead to uncovering more accurate biological insights, as bulk proteomics functions only as proxy by averaging thousands of different cells and cell types. Since the first introduction of single-cell proteomics (SCP) methods in 2018^1,2^ the field made enormous advances, now enabling deep proteome coverage of >5000 proteins from a single cell with high quantitative performance^3,4^. This progress was largely driven by technological innovations like the new generations of sensitive mass spectrometer instruments such as the Orbitrap Astral and timsTOF Ultra. Two complementary quantification strategies dominate this progress: isobaric labeling using tandem mass tags (TMT) and label-free quantification (LFQ) using data-independent acquisition (DIA). DIA-LFQ is gaining more and more traction especially in the SCP field due to its superior depth and data completeness across replicates compared to TMT-based approaches which have conventionally relied on data-dependent acquisition (DDA)^5^. However, TMT remains attractive because it enables high sample throughput via multiplexing of up to 35 samples (or single cells) in a single MS run, with throughput being crucial for statistical power in single cell studies. On the contrary, isobaric labeling introduces caveats, such as ratio compression^6–8^, higher sample preparation costs and a more time-consuming sample preparation. As TMT data is recorded in DDA mode, this approach is limited by its stochastic precursor selection, which can lead to missing values across runs eventually resulting in lower reproducibility^9,10^. Furthermore, low-abundant peptides may be overlooked, as intensity thresholds favor the sampling of highly abundant peptides^11,12^. These limitations are even more pronounced for limited input, which is especially true in SCP. In contrast, DIA-LFQ provides comprehensive and reproducible peptide detection across samples by fragmenting all precursor ions within a defined *m/z* isolation window, avoiding the stochastic nature of DDA. Therefore, a desirable synergistic effect is anticipated when combining both DIA and TMT-based multiplexing due to their complementary strengths. The general feasibility of DIA-TMT was already shown by Ctortecka and colleagues, but without peptide identifications connected to their quantitative profiles^13^. The integration of TMT labeling with DIA presents technical challenges, particularly in terms of interference correction, quantification accuracy and data analysis complexity. A major concern in this regard is co-isolation interference, where co-fragmented ions from different peptides contribute to the same fragment spectra, distorting TMT reporter ion (RI) signals. This leads to increased ratio compression and compromises the accuracy of quantitative measurements. Thus, a general recommendation for acquiring TMT data is to reduce the isolation window width (≤ 1 *m/z*), already minimizing the co-isolation of precursors, alleviating true ratio distortion between samples or single cells^7,14^. An alternative approach to circumvent RI-based interference altogether is quantification via TMT complementary ions (TMTc), which are generated as a byproduct of RI formation and retain the precursor’s isotope envelope^15^. Because TMTc ions are typically of higher *m/z* and lower relative interference than reporter ions, they can, in principle, offer improved quantitative accuracy at the cost of additional data analysis complexity. When using isolation windows ≤ 1 *m/z* in DIA, the cycle time increases dramatically, limiting its applicability to fast state-of-the-art instruments such as the Orbitrap Astral mass spectrometer with high sampling rate and parallelized ion packet handling^16^. The Astral platform combines high-resolution Orbitrap detection with an ultra-fast Astral analyzer, enabling rapid and parallelized acquisition of high-resolution MS/MS spectra. These features are particularly well suited for DIA-TMT workflows, which demand both high scan speed and precision to resolve multiplexed TMT RIs and complex fragment ion patterns^16,17^.

Despite this promise, a systematic, identification-linked benchmarking of DIA-TMT across a broad range of acquisition parameters has not yet been performed on state-of-the-art instrumentation such as the Orbitrap Astral mass spectrometer. Such an evaluation is needed to connect quantitative accuracy, interference levels and proteome coverage to specific instrument settings. To address this gap, we devised a two-proteome interference model consisting of human and yeast digests. This model allowed controlled assessment of interference effects and quantification accuracy by introducing known ratios and complexity. We tested a range of MS acquisition strategies, varying in isolation windows, injection time (IT) and input amount to identify optimal conditions for DIA-TMT analysis. Using our in-house Python tool, **R**eporter Ion **E**xtractor and **A**nnotation **D**irector (READ), we quantified TMT RIs and calculated co-isolation purity to enable systematic quality control across acquisition methods, offering a toolkit for TMT-based quantification independent of the search engine’s native reporter-ion support. We also included an analysis of TMTc ions instead of TMT RIs to alleviate reduced quantitative accuracy caused by precursor co-isolation.

## Results and discussion

### Experimental design and data quality of a two-proteome interference model

Two-proteome interference models are widely recognized experimental designs for evaluating the quantitative performance of LC-MS/MS methods^3,18–20^. Two-proteome mixtures, as employed here, are particularly well suited to benchmark TMT-based workflows, where ratio compression represents a major challenge for accurate quantification. We combined human and yeast proteomes at three yeast spike-in levels (H1:Y0.1, H1:Y0.2 and H1:Y1), corresponding to yeast fold changes (FC) of 1:2, 1:5 and 1:10 between the mixes, distributed across TMTpro 18-plex channels with three intra-sample replicates each (Figure 1B). We also included empty channels to investigate the signal bleeding from neighboring channels. This design was used to benchmark a panel of DDA and DIA acquisition methods on the Orbitrap Astral mass spectrometer, varying isolation window size, MS2 IT and injection amount (Figure 1C). Before comparing methods systematically, we first assessed data quality for a representative DDA method (0.5 *m/z* isolation window, 8 ms IT, 54 ng input). The distribution of summed TMT RI at the protein level showed high consistency across the intra-sample replicates, although channel 135N was an exception: the reduced intensity of 1.7E8 in comparison to 2.1E8 for 134N and 134C is consistent with lower TMT labeling efficiency in that channel (Figure 1D). Furthermore, we observed low signal bleeding into the empty channels (e.g., summed RI intensity of 6.8E5 in 127C), which increased to a summed RI intensity of 1.1E7 in 133C due to higher inputs in the neighboring channels. As a second quality control, we evaluated quantitative accuracy by comparing human protein ratios between the HeLa control and Mix1 channels using a Bland-Altman plot. Because we kept the human background constant in all peptide containing channels, the theoretical FC is 1 (Figure 1E; same method as in Figure 1D). Across 4217 quantified human proteins, the mean measured FC was 0.9, indicating good agreement with the expected ratio and only a minor deviation. Together, these results confirm that the experimental design and data quality are suitable for the systematic method comparison presented below.

**Figure 1:**
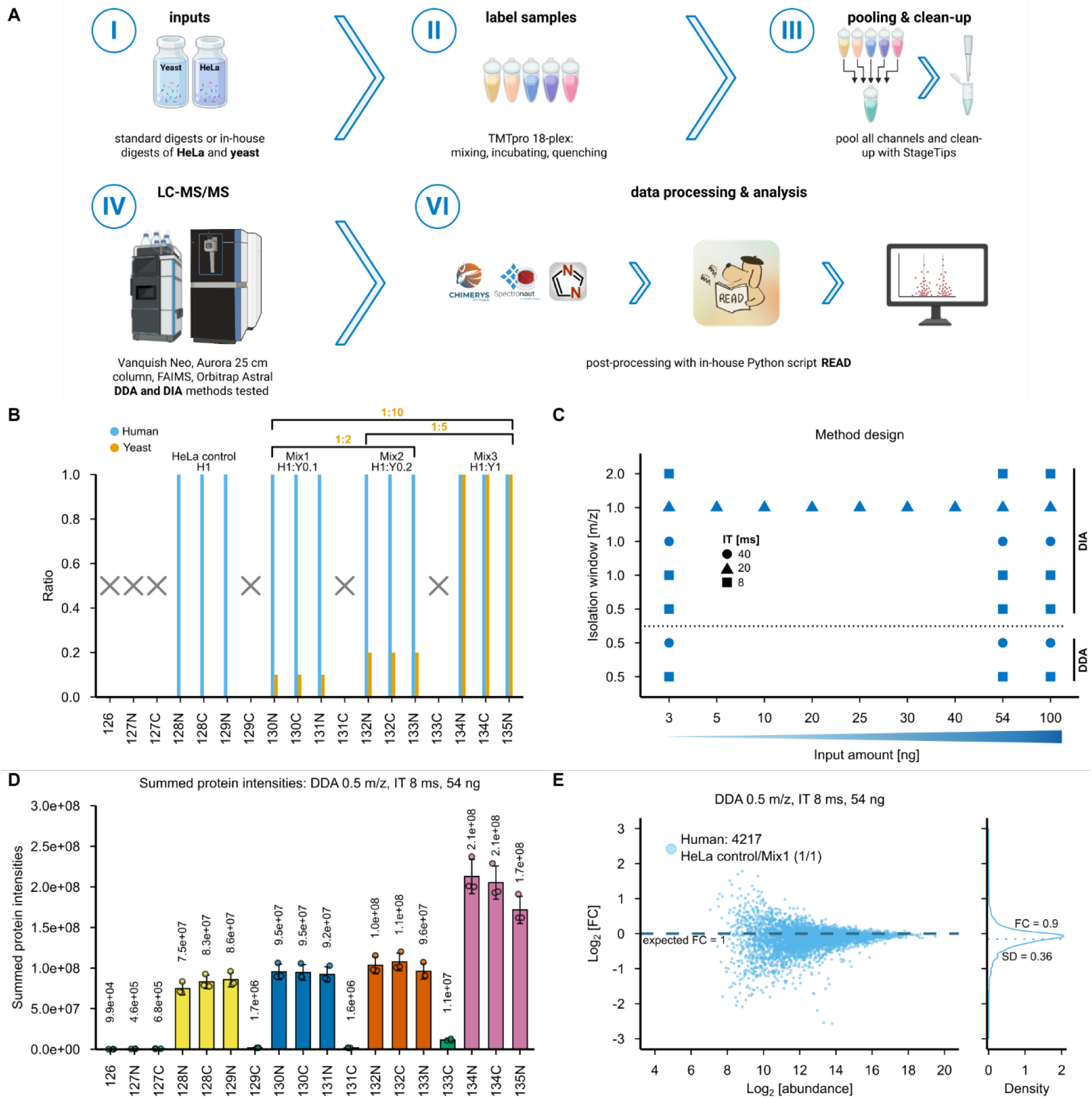
Experimental overview and first quality control of the data. (**A**) Schematic overview of the experimental and data analysis workflow: human and yeast tryptic peptides are combined at defined ratios, labeled with TMTpro 18-plex reagents, pooled, and desalted via StageTips prior to LC-MS/MS analysis on a Vanquish Neo coupled to an Orbitrap Astral mass spectrometer (25 cm Aurora column, FAIMS) using both DDA and DIA methods. Raw data were processed with CHIMERYS, Spectronaut, or DIA-NN, followed by reporter ion (RI) quantification and co-isolation assessment using the in-house Python tool READ. (**B**) The TMTpro 18-plex set with six empty channels, three intra-sample replicates for the pure human channels (HeLa control) and three intra-sample replicates each for the two-proteome mixes containing increasing amounts of spiked-in yeast peptides to a constant human peptide background (Mix1-Mix3). (**C**) Overview of the tested DIA and DDA MS method settings on the Orbitrap Astral mass spectrometer with different isolation windows, MS2 injection times (IT) and injection amounts. (**D**) Bar plot of TMT RI intensities for a representative DDA method (0.5 *m/*z isolation window, MS2 IT 8 ms, injection amount of 54 ng; bars indicate average values, error bars show their standard deviation, n = 3 technical replicates). (**E**) Representative Bland-Altman plot from one technical replicate of (**C**) showing measured log2 fold changes (FC) for human proteins between HeLa control and Mix1, where the theoretical FC is 1 (dashed line). Density plot on the right shows the distribution of measured log2 FC with the mean indicated as a dashed line (n = 3 intra-sample replicates).

### Input amount and acquisition parameters govern co-isolation and protein identification

To investigate how sample input amount affects protein quantification, we performed an input titration using a selected DIA method (1 *m/z* isolation window, MS2 IT 40 ms) across a range of inputs from 3 to 100 ng (Figure 2A). This range was chosen to reflect biologically relevant SCP setups: 3 ng corresponds to the estimated protein content of 12 multiplexed single cells at 250 pg each, while 54 ng corresponds to TMT-SCP setups comprising a carrier channel of 200 single cells combined with an empty channel and with 16 actual single cells across the remaining TMTpro channels corresponding to the cell number used in the SCoPE-MS protocol and adopted by others^2,21^. The number of quantified proteins increased with input amount, from 5609 (3 ng) to 9702 (100 ng), however the increase leveled off at higher loads. Because co-isolation purity is calculated per precursor at the MS1 isolation step but is only accessible for precursors that were subsequently identified, all purity distributions and threshold-based percentages reported below refer to identified PSMs (as reported by CHIMERYS in Proteome Discoverer for both DDA and DIA), rather than the full population of isolated precursors. This trend was mirrored in the co-isolation purity-distributions (Figure 2C): higher input amounts shifted the distribution to lower purity values, with the fraction of PSMs above 70% purity decreasing from 24.3% at 3 ng to 18.7% at 100 ng, consistent with a higher density of co-eluting precursors at increased input amount. Applying stringent filtering (co-isolation purity ≥ 70%, TMT RI signal-to-noise ratio (S/N) ≥ 20, TMT RI resolution ≥ 45,000) substantially reduced the number of quantified proteins across all inputs (e.g., from 5609 to 602, a 10.7% retention rate, compared to 9702 to 4214 at 100 ng, a 43.3% retention rate), most notably at low input amounts, where the minimum S/N requirement was fulfilled less frequently. To select appropriate filter thresholds, we evaluated a representative DDA (0.5 *m/z*, 8 ms IT) and DIA (1.0 *m/z*, 8 ms IT) method at increasing S/N cut-offs (3-80), in combination with the fixed purity (≥70%) and resolution (≥45,000) thresholds (Supplementary Figure 2). Increasing S/N stringency progressively reduced the number of quantified proteins, more markedly for DIA than DDA (Supplementary Figure 2A), while yielding modest improvements in quantitative accuracy (Supplementary Figure 2C). Coefficient of variation (CV) distribution showed a non-monotonic trend, remaining high directly after purity filtering, transiently decreasing at intermediate S/N thresholds, and recovering at higher thresholds (Supplementary Figure 2B), potentially reflecting more homogeneous RI quantification once low-confidence noise-dominated signals are excluded. Given the limited additional gain in accuracy at very high S/N thresholds relative to the associated loss in proteome depth, we selected S/N ≥ 20 as a pragmatic compromise for the remaining analyses.

**Figure 2:**
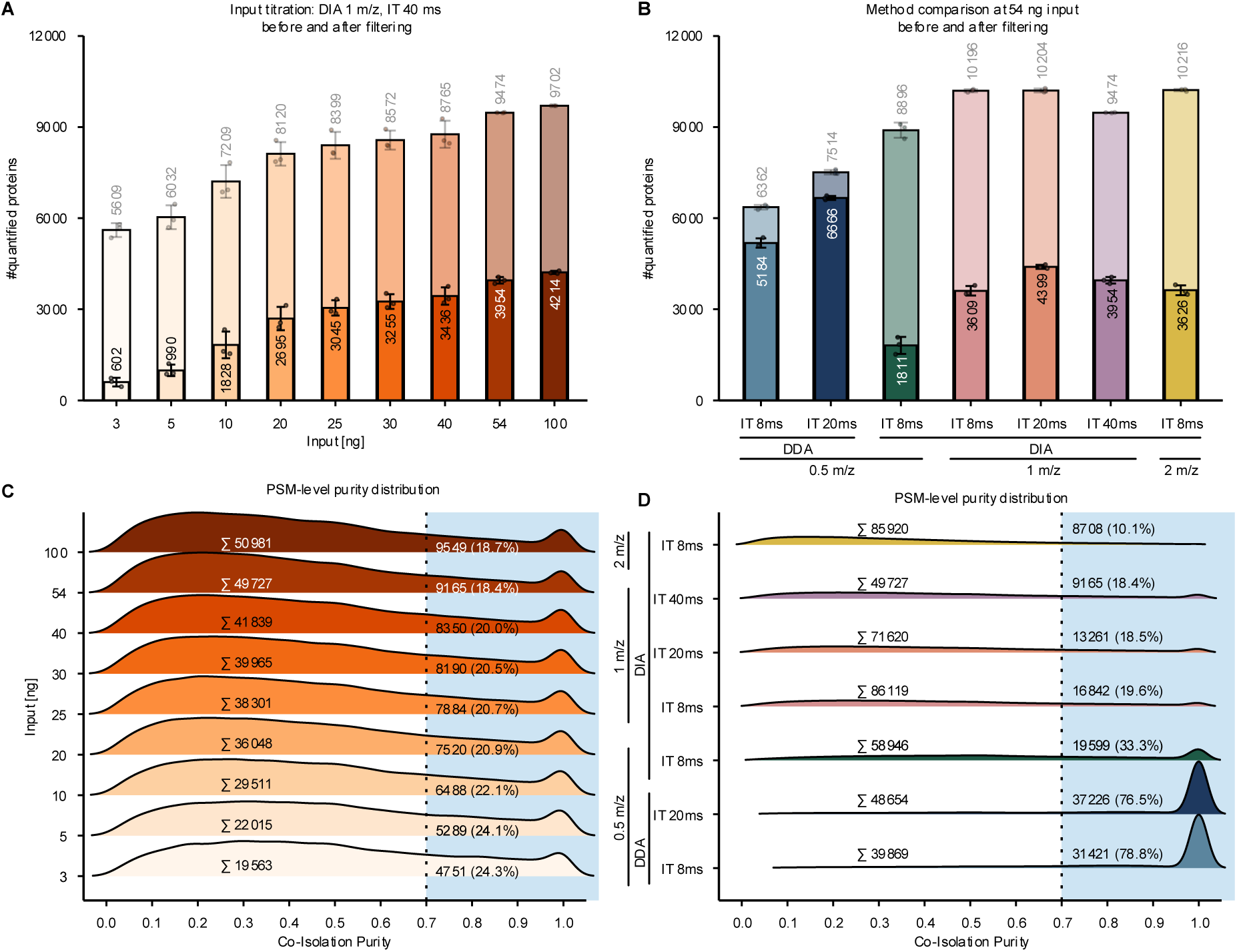
Input titration and acquisition method benchmark. (**A**, **B**) Bar chart showing the number of quantified proteins with or without filtering for co-isolation purity ≥ 70%, TMT reporter ion (RI) signal-to-noise ratio ≥ 20 and TMT RI resolution ≥ 45,000 when acquiring data at an isolation window of 1 *m/z* and 40 ms MS2 injection time (IT) with varying inputs as indicated (**A**) or at 54 ng input with varying isolation windows and injection times as indicated (**B**; bars indicate average values, error bars their standard deviation, n = 3 technical replicates). (**C**, **D**) Density distribution for all PSMs (identified by CHIMERYS in Proteome Discoverer) at given co-isolation purity in dependence of sample input (**C**) or acquisition method (**D**); n = 3 technical replicates.

We next compared DDA and DIA acquisition methods at a fixed input of 54 ng, varying isolation window size and MS2 IT (Figure 2B). While DIA methods initially quantify more proteins than DDA methods, this trend is reversed as soon as the above-mentioned filter criteria are set. Both MS2 IT and isolation window size have an influence on the number of quantified proteins. Within DIA methods using the same isolation window, increasing MS2 IT did not change co-isolation purity distribution per PSM, as the fraction of PSMs passing the 70% purity threshold remained largely unchanged across IT settings (Figure 2D). This was expected, since purity is set at the MS1 isolation step and is therefore independent of MS2 IT. Instead, longer MS2 ITs reduced the total number of precursors triggered for MS2 within the fixed gradient time, as fewer MS2 scans fit into the same acquisition window. Since the proportion of high-purity PSMs stayed constant, this reduced sampling depth directly translated into fewer purity-passing PSMs and, consequently, fewer quantified proteins at longer ITs. At the same time, longer MS2 ITs are expected to increase the TMT RI S/N through longer ion accumulation, which can reduce the number of missing RI values in successfully quantified PSMs. Thus, longer MS2 IT introduces a trade-off in DIA-TMT workflows: while sampling depth and the number of quantified proteins decrease, the quality of RI quantification per PSM is expected to improve.

Isolation window width had a stronger effect on co-isolation than MS2 IT. Broader isolation windows led to a higher number of co-isolated precursors, both in absolute terms and as a fraction of all detected PSMs, consistent with the increased probability of capturing multiple co-eluting species within a wider *m/z* range (Figure 2C, D). This effect was particularly pronounced for DIA methods, where isolation windows are fixed and applied irrespective of the underlying precursor distribution, making wider DIA windows especially susceptible to co-isolation. Notably, when DIA was run with a narrow 0.5 *m/z* isolation window, identical to the isolation window used in DDA, the number of quantified proteins was nevertheless lower than for DIA at 1 *m/z* or DDA at 0.5 *m/z* (Figure 2B), despite the narrower window improving co-isolation purity (Figure 2D). In DDA, the isolation window is centered around the precursor while in DIA the precursors can lie anywhere in the isolation window, thereby increasing the likelihood of co-isolation.

This suggests that narrowing the DIA isolation window trades off co-isolation purity against the number of windows required to cover the same precursor mass range, thereby increasing overall DIA cycle time. A longer cycle time reduces the number of complete cycles sampled across the chromatographic peak, which can impair confident peptide identification in DIA. Additionally, narrower isolation windows capture fewer ions per scan, resulting in lower RI intensities, noisier RI signals, and ultimately reduced protein quantification quality. In comparison, DDA methods retained the highest number and proportion of PSMs above the 70% purity threshold (76.5-78.8% for 0.5 *m/z* DDA vs. 10.1-33.3% across DIA isolation windows) (Figure 2B, D). This is further supported by the fact that DDA acquisition on Thermo instruments inherently incorporates an isotope envelope-based selection step, the Precursor Fit filter, which was set to a minimum of 70% in this study and compares the observed isotopic pattern of a candidate precursor to its theoretical envelope, preferentially triggering MS2 on precursors with a good fit^22^, i.e. low co-isolation. This is consistent with the high measured purities observed for DDA above. Nevertheless, a fraction of DDA PSMs (21.2–23.5%) still fell below the 70% threshold despite this real-time filter. This discrepancy likely reflects differences between the real-time purity estimate used by the instrument at the moment of precursor selection, and the purity subsequently calculated from the processed experimental data. In contrast, DIA methods isolate and fragment all ions within a predefined window irrespective of isotope fit, making them inherently more susceptible to co-isolation interference, particularly at wider isolation window settings.

Taken together, these results show that input amount, isolation window width and MS2 IT determine the extent of co-isolation interference and consequently the number of quantifiable proteins. This highlights the need to carefully balance sensitivity and selectivity when designing TMT-based DIA methods.

### Ratio compression and its dependence on acquisition parameters and analysis software

With our interference models we were also able to reveal potential issues with ratio compression, an artifact of TMT-based quantification arising from co-isolation of interfering ions^7^. We compared quantitative accuracy and precision between representative DDA and DIA methods using Bland-Altman plots of human and yeast proteins between Mix2 and Mix1 (Figure 3A, B). For yeast proteins, the theoretical FC of 2 was substantially underestimated in both methods (measured mean FC = 1.29 for DDA and 1.23 for DIA), while human protein ratios, with an expected FC of 1, showed only minor deviation (measured FC = 1.05 and 1.02, respectively). This asymmetric bias is characteristic of ratio compression. Since the human background is present at constant high abundance across all channels, co-isolated human precursor signal dilutes the yeast-specific RI signal, pulling the measured yeast ratio toward 1. As human ratios are close to 1 already, compression toward 1 has little visible effect on their accuracy. Notably, DDA and DIA performed comparably poorly in this regard, indicating that ratio compression here is driven primarily by the complexity and lack of fractionation of the sample, rather than by the acquisition mode itself. This is consistent with findings from He *et al*.^19^, who reported compressed ratios for DDA-TMT on the Orbitrap Astral mass spectrometer even at moderate spike-in ratios (e.g., a theoretical FC of 2 was measured as 1.59; Supplementary Figure 11b in^19^). Notably, their samples were fractionated prior to MS analysis, which is expected to reduce sample complexity, thereby alleviating co-isolation and compression. Despite this, their fractionated samples still showed substantial compression, underscoring that even with reduced sample complexity, ratio compression remains a persistent challenge for TMT RI-based quantification. This suggests that ratio compression is likely to be even more pronounced in our unfractionated, low-input two-proteome model, which mimics the sample constraints relevant to SCP, where fractionation is not feasible due to the limited input amount. The magnitude of this distortion scaled with the true FC: at higher yeast spike-in ratios (Mix3/Mix2 and Mix3/Mix1 with expected FC of 5 and 10, respectively), yeast ratios compression became more pronounced (measured FC = 2.86–3.37 and 3.48–4.31, respectively) and, notably human protein ratios (expected FC = 1) became clearly inflated (measured FC = 1.53-1.68) (Supplementary Figure 3). This mirrors previous findings that co-isolation of a differentially abundant interfering species can inflate the measured ratio of an otherwise constant background protein, as its RI signal becomes increasingly contaminated by the diverging interferant^8,23^.

**Figure 3:**
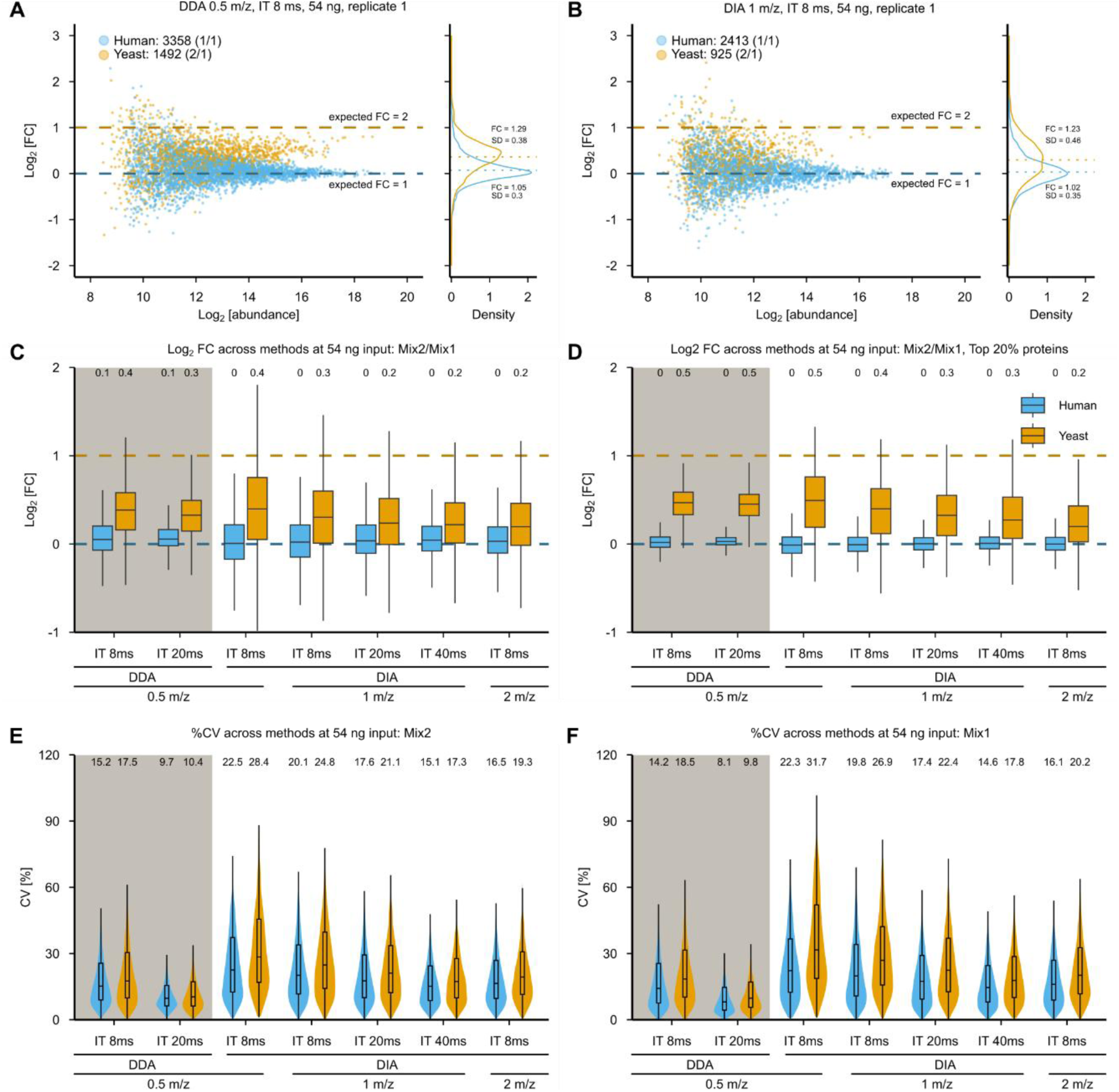
Accuracy and precision of quantification in a two-proteome mixture across DDA and DIA acquisition methods. The shown data was filtered for co-isolation purity ≥ 70%, TMT reporter ion signal-to-noise ratio ≥ 20 and TMT reporter ion resolution ≥ 45,000. (**A**, **B**) Representative Bland-Altman plot from one technical replicate of a DDA (0.5 *m/*z, MS2 IT 8 ms, 54 ng) and a DIA (1 *m/*z, MS2 IT 8 ms, 54 ng) measurement showing measured log2 fold changes (FC) for human and yeast proteins between Mix2 and Mix1, where the theoretical log2 FC is 0 for human (dashed blue line) and 1 for yeast proteins (dashed orange line). Density plots on the right show the distribution of measured log2 FC with the mean indicated as a dashed line (n = 3 intra-sample replicates). (**C**) Box plots showing the log2 FC for human (blue) and yeast proteins (orange) between Mix2 and Mix1 (as in **A** and **B**) across indicated DIA and DDA methods with an input of 54 ng. The horizontal center line represents the median, the lower and upper hinges correspond to the first and third quartiles (25^th^ and 75^th^ percentiles, defining the interquartile range (IQR)) and the whiskers extend to the furthest data points within 1.5x IQR from the hinges. Outliers are omitted from the visualization for clarity. Values above each box represent the corresponding median. Horizontal dashed lines indicate theoretical/expected log2 FC (0 for human and 1 for yeast proteins) (n = 3 technical replicates). (**D**) As in (**C**) but showing only the log2 FC-distribution of the top 20% for each comparison/organism. (**E**, **F**) Violin plots show the distribution of coefficients of variation (CV [%]) for human (blue) and yeast (orange) proteins across the indicated methods for Mix2 (E) and Mix1 (F). Violin width reflects the relative density of data points at a given CV value. Overlaid boxplots indicate the median (center line) and IQR (25^th^-75^th^ percentiles, box limits); outliers are omitted for clarity. Numbers above each violin show the respective median CV (n = 3 technical replicates).

Across the panel of DDA and DIA methods benchmarked in Figure 2B, quantitative accuracy for the Mix2/Mix1 comparison varied only modestly (Figure 3C): narrower isolation windows and shorter MS2 IT were associated with improved accuracy (e.g., median log2 yeast FC of 0.4 for the narrowest DIA condition (0.5 *m/z*) versus 0.2 for the broadest DIA condition (2 *m/z*); human medians remained at 0-0.1 across all methods), consistent with the reduced co-isolation under these conditions (Figure 2D). However, the magnitude of these differences was very small relative to the overall compression observed across all methods, reinforcing that the co-isolation interference and not the acquisition parameters are the main drivers of ratio compression in this dataset. When restricting the analysis to the top 20% most abundant proteins for each organism (Figure 3D), accuracy improved consistently across most methods and narrowed the spread of the data (e.g., yeast median log2 FC increased toward the expected value of 1 across most conditions, reaching 0.5 for DIA 0.5 *m/z*, while human medians remained near 0). This is expected, as highly abundant precursors are less susceptible to interference from lower-abundance co-isolating species.

In contrast to accuracy, quantification precision showed a clearer dependence on acquisition parameters (Figure 3E, F). DIA methods generally exhibited higher CVs than DDA methods, and precision improved with wider isolation windows and longer MS2 ITs (e.g., median yeast CV decreased from 24.8% at 8 ms IT to 17.3% at 40 ms IT for DIA with an isolation width of 1 *m/z*; DDA 20 ms IT showed the lowest overall CVs, 9.7%/10.4% for human/yeast; Figure 3E). This apparent trade-off mirrors the sampling-depth/signal-quality relationship described above (Figure 2): wider windows and longer ITs increase the number of ions collected per precursor, improving reporter ion S/N and thus quantitative precision, even though the same conditions increase co-isolation and therefore compromise accuracy. Together, these results indicate that accuracy and precision in DIA-TMT workflows are governed by partially opposing requirements.

We also evaluated whether search software choice affects TMT RI-based quantification by comparing Proteome Discoverer, DIA-NN and Spectronaut using the same DIA method (1 *m/z* isolation window, MS2 IT 8 ms, 54 ng; Supplementary Figure 4A). DIA-NN quantified the highest number of proteins for both before and after filtering (11,091 and 4,315, respectively) followed by Proteome Discoverer (10,196 and 3,609) and Spectronaut (9,420 and 3,416), indicating differences in identification sensitivity between the three tools. Despite these differences in protein numbers, quantitative accuracy for the Mix2/Mix1 comparison was comparable across all three software packages (Supplementary Figure 4B): yeast ratios were compressed similarly (median log2 FC of 0.2-0.3), and human ratios remained at the expected log2 FC of 0 in all cases. Precision assessed via CV distributions for Mix2 and Mix1 (Supplementary Figure 4C and D, respectively) was likewise similar across software tools, with median CVs ranging from 18.2-20.1% for human and 22.6-26.9% for yeast proteins. Together, these results indicate that the choice of search software has only a minor influence on quantitative accuracy and precision in our TMT RI-based DIA-TMT workflow, confirming that the ratio compression described above is not specific to a single analysis pipeline.

### A single-proteome dynamic range benchmark confirms high quantitative performance across DDA and DIA

To assess TMT RI-based quantification performance under conditions with minimal co-isolation interference, we designed a loading scheme comprising four relative human protein input levels: H0, H1, H5 and H10, corresponding to relative loading ratios of 0x, 1x, 5x and 10x. These groups were distributed across the TMTpro 18-plex with intervening empty channels to monitor signal bleeding (Figure 4A). Unlike the two-proteome interference model, this design avoids mixing species, thereby minimizing co-isolation-driven interference. For a representative DDA method (0.5 *m/z*, MS2 IT 20 ms, 54 ng), summed RI intensities scaled consistently with the intended loading ratios across replicates. Signal bleeding into empty channels remained low relative to the neighboring loaded channels, confirming the suitability of this design.

**Figure 4:**
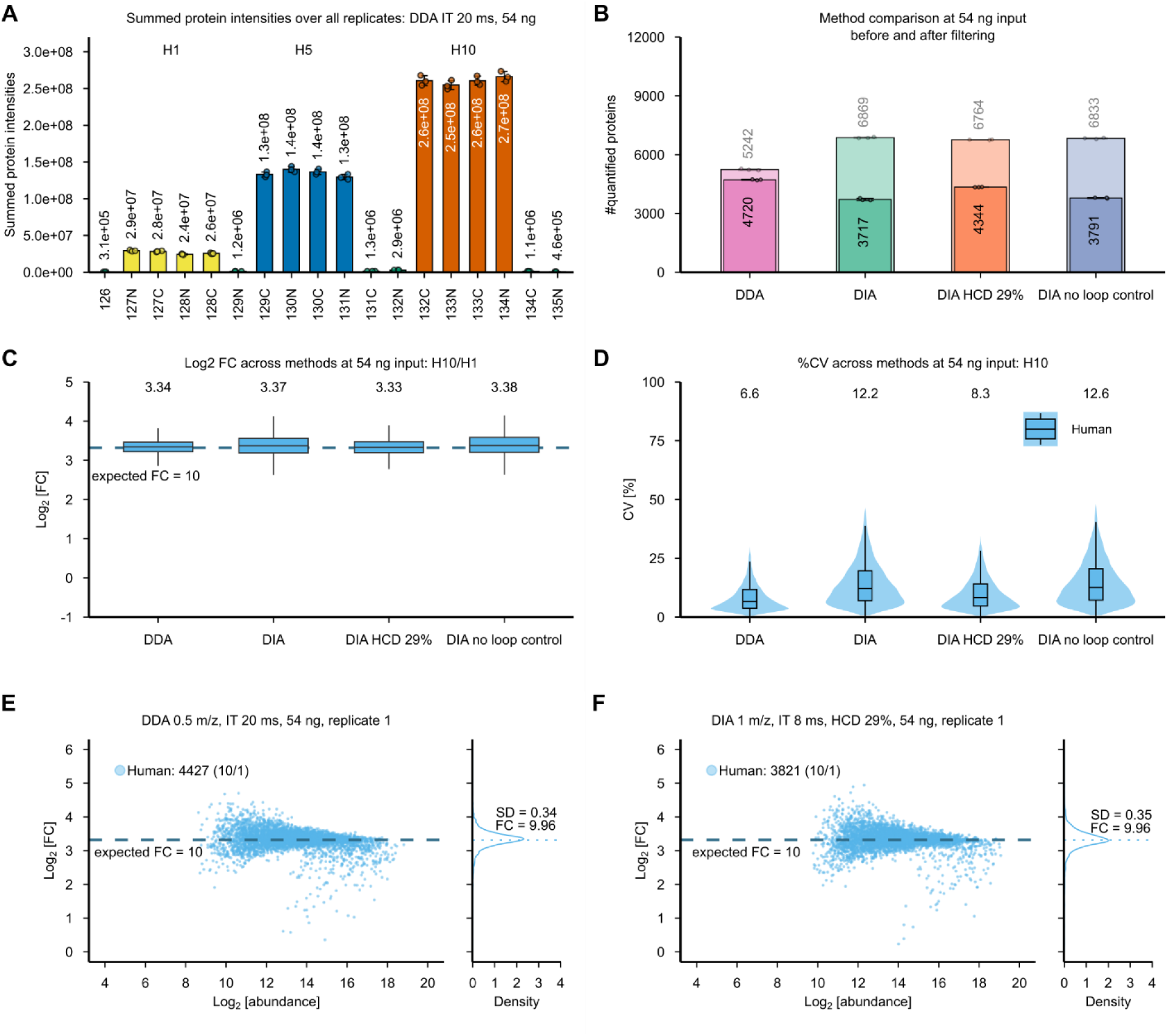
Benchmarking TMT reporter ion (RI)-based quantification across DDA and DIA acquisition methods. The data shown in B, C, D, E and F was filtered for co-isolation purity ≥ 70%, TMT reporter ion signal-to-noise ratio ≥ 20 and TMT reporter ion resolution ≥ 45,000. (**A**) Summed protein intensities across all 18 TMTpro channels for a representative DDA method (0.5 *m/z*, MS2 IT 20 ms, 54 ng), grouped into four loading levels: H0 (green), H1 (yellow), H5 (blue) and H10 (orange). Values above each bar indicate summed RI intensities; dots represent individual replicate values, bars indicate average values, error bars their standard deviation (n = 3 technical replicates). (**B**) Number of quantified proteins at 54 ng input across the indicated methods: DDA (as in **A**), DIA (1 *m/z*, MS2 IT 8 ms, HCD 25%, loop control 0.6s), DIA HCD 29% (1 *m/z*, MS2 IT 8 ms, HCD 29%, loop control 0.6s) and DIA (1 *m/z*, MS2 IT 8 ms, HCD 25%, loop control disabled). The number of quantified proteins is given before (light, top values) and after (solid, bottom values) filtering (co-isolation purity ≥ 70%, TMT RI signal-to-noise ratio ≥ 20 and TMT RI resolution ≥ 45,000; n = 3 technical replicates). (**C**) Box plots showing the log2 fold change (FC) for human proteins between H10 and H1 across indicated DIA and DDA methods with an input of 54 ng. The horizontal center line represents the median, the lower and upper hinges correspond to the first and third quartiles (25^th^ and 75^th^ percentiles, defining the interquartile range (IQR)) and the whiskers extend to the furthest data points within 1.5x IQR from the hinges. Outliers are omitted from the visualization for clarity. Values above each box represent the corresponding median. Horizontal dashed lines indicate theoretical/expected log2 FC of 3.32 (n = 3 technical replicates). (**D**) Violin plots show the distribution of coefficients of variation (CV [%]) for human proteins across the indicated methods for H10. Violin width reflects the relative density of data points at a given CV value. Overlaid boxplots indicate the median (center line) and IQR (25^th^-75^th^ percentiles, box limits); outliers are omitted for clarity. Numbers above each violin show the respective median CV (n = 3 technical replicates). (**E**, **F**) Representative Bland-Altman plot from one technical replicate of a DDA (E; 0.5 *m/*z, MS2 IT 20 ms, 54 ng) and a DIA HCD 29% (F; 1 *m/z*, MS2 IT 8 ms, HCD 29%, loop control 0.6s) measurement showing measured log2 FC for human proteins between H10 and H1, where the theoretical log2 FC is 3.32 for human (dashed blue line). Density plots on the right show the distribution of measured log2 FC with the mean indicated as a dashed line (n = 3 intra-sample replicates).

We compared this DDA method against three DIA method variants at the same 54 ng input (Figure 4B): a standard DIA method (1 *m/z*, MS2 IT 8 ms, higher-energy collisional dissociation (HCD) 25%, loop control 0.6 s), DIA with increased fragmentation energy (HCD 29%), and DIA without loop control. We reasoned that the standard HCD energy, while optimized for peptide backbone fragmentation in DIA-LFQ, may under-fragment TMT RI, thereby limiting RI intensity and S/N. Thompson *et al*. reported that normalized collision energies (NCEs) of 30-32% are optimal for TMTpro RI generation^24^, while Wang *et al*. found that HCD energies at 29% are optimal for generation of complementary ions^25^. Taken together, these findings suggest that higher NCEs than those used in our default method are generally advantageous for RI generation. In line with this literature, we therefore tested whether increasing HCD energy to 29% improves RI signal in our DIA workflow. We also tested disabling loop control, which normally caps the maximum cycle time to maintain a defined duty cycle. This was expected to increase MS sampling depth and potentially the number of quantified proteins, but at the cost of the duty-cycle constraint that otherwise balances ion accumulation time against scan speed. Before filtering, all three DIA methods quantified more proteins than DDA (6764–6869 vs. 5242). After applying our quality filters (co-isolation purity ≥ 70%, TMT RI S/N ≥ 20, TMT RI resolution ≥ 45,000), DDA retained the highest number of quantified proteins (4720), followed by DIA HCD 29% (4344), DIA no loop control (3791) and standard DIA (3717). This indicates that increased HCD energy can partially narrow the gap to DDA.

Quantitative accuracy for the H10/H1 comparison (theoretical log2 FC = 3.32) was high and comparable across all four methods (Figure 4C), with median log2 FC values ranging narrowly from 3.33 to 3.38. This demonstrates that, in the absence of substantial co-isolation interference, both DDA and DIA achieve accurate TMT RI-based quantification regardless of acquisition mode, in clear contrast to the pronounced ratio compression observed in Figure 3.

Quantification precision was likewise high overall in the H10 group but varied somewhat between methods (Figure 4D). DDA showed the lowest median CV (6.6%), while DIA with no loop control showed the highest (12.6%). Increasing HCD energy to 29% improved DIA precision (8.3%), consistent with our expectation that higher fragmentation energy increases RI signal quality. Disabling loop control worsened precision slightly compared to standard DIA (12.2%).

Representative Bland-Altman plots for DDA and DIA HCD 29% (Figure 4E, F) illustrate the comparable accuracy (measured FC = 9.96 for both) and similar spread (SD = 0.34 vs. 0.35) achieved by both acquisition modes. Together, these results confirm that when co-isolation interference is minimized, TMT RI-based quantification is accurate and precise across both DDA and DIA acquisition modes, and that RI signal quality in DIA can be further improved through targeted parameter adjustments such as increased HCD energy.

### TMTc-based quantification trades sensitivity for reduced co-isolation susceptibility relative to TMT RI

TMTc-based quantification has been proposed as an alternative to conventional TMT RI-based quantification, since it is inherently resistant to co-isolation interference^15,26^. Rather than relying on the low-mass RI, which are indistinguishable across co-isolated peptides, the TMTc approach exploits the heavier complementary ions retained on the peptide backbone after HCD-induced loss of the reporter group and a neutral carbon monoxide. These complementary ions are detected at a substantially higher mass range, up to 2000 *m/z* and beyond. We adapted this strategy for our DIA workflow, selecting the TMTpro channels 126, 131N and 135N, consistent with Wang *et al*.^25^. These channels produce complementary ions spaced 4 Da apart, sufficiently distant to minimize isotopic envelope overlap and avoid the need for complex deconvolution. In theory, the TMTpro 18-plex could yield up to nine distinguishable complementary ion masses, since channel pairs collapse on the same complementary ion mass after loss of the reporter group. In practice, resolving these closely spaced complementary ions would require a resolution up to or even higher than 600,000^27^, which is not achievable with current Orbitrap instrumentation, including the Astral platform used here.

Our implementation differs from Wang *et al*.^25^ in several methodological aspects. We employed an Orbitrap Astral mass spectrometer instead of an Orbitrap Exploris 480. We also used a narrower 8 *m/z* isolation window compared to their 16 *m/z* window for DIA acquisition. We evaluated TMTc quantification across four methods: DDA, standard DIA (60 min gradient), a higher-mass DIA variant (HM-DIA, 60 min) and a 120 min gradient method (DIA120). We further aimed for working at low input amounts (50 and 100 ng), consistent with our preceding DIA-TMT RI benchmarking, targeting low-input and SCP-relevant conditions.

As a quality control step, we first confirmed that the measured TMTc signal reflects the intended experimental design. Figure 5A shows the summed protein intensities derived from TMTc ion quantification across the three channels for a representative method (DIA120 with 50 ng input), after subtracting a species-specific noise (median + one SD of the empty channel 126). Human protein intensities were comparable for 131N and 135N, while yeast protein intensities showed a ratio close to the expected FC of 5, confirming that the measured signal accurately reflects the intended label design.

**Figure 5:**
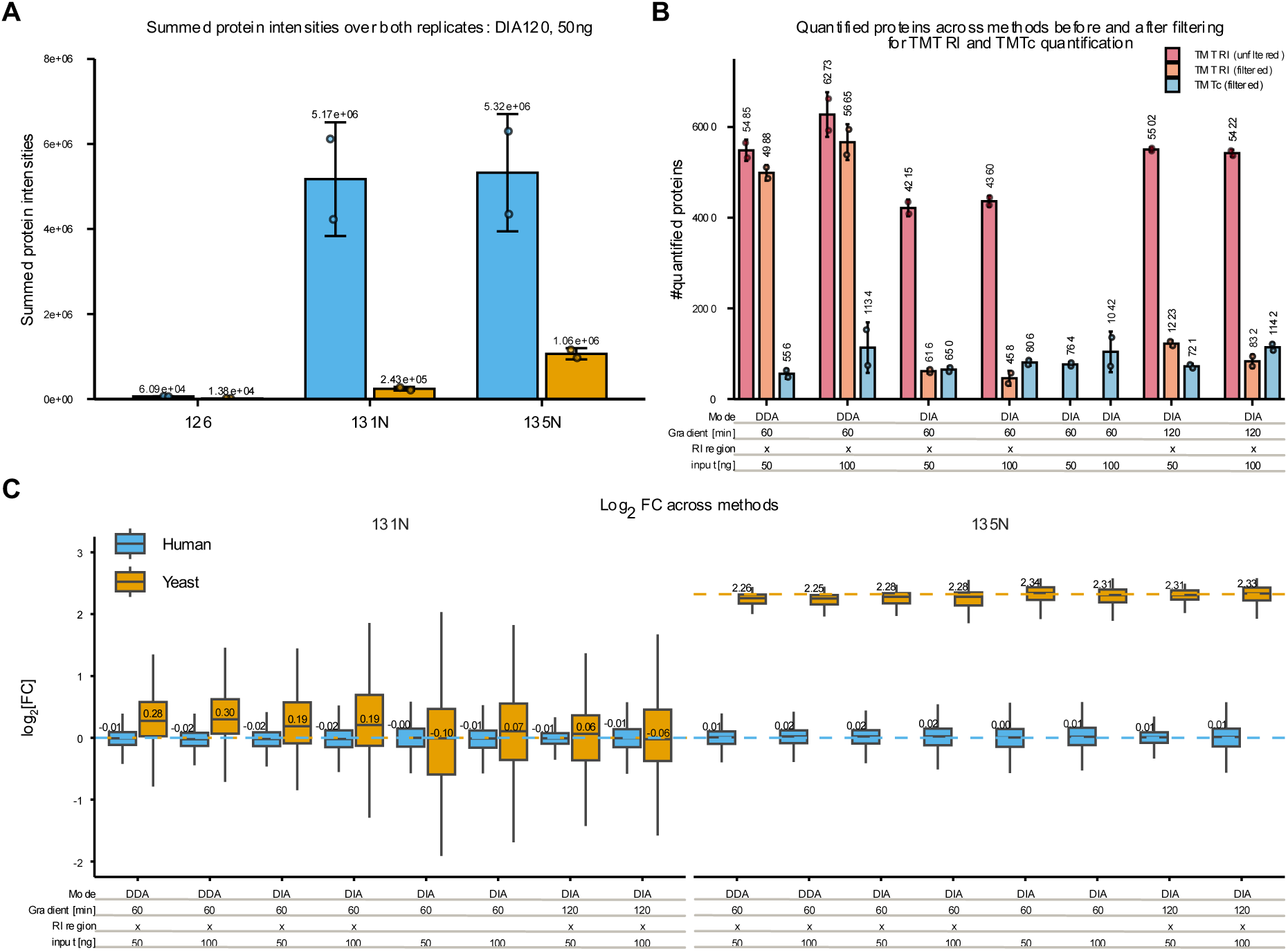
Benchmarking TMT complementary ion (TMTc)-based quantification against TMT reporter ion (RI)-based quantification across DDA and DIA acquisition methods. (**A**) Summed protein TMTc intensities across TMTpro channels 126, 131N and 135N after subtracting the species-specific noise threshold (median + one standard deviation (SD) of the empty 126 channel) from all three channels, averaged over both replicates for a representative DIA120 method (50 ng), shown for yeast (orange) and human (blue) proteins. Channel 126 (empty for both species) is shown for reference; any apparent residual signal reflects background noise exceeding its own threshold by chance, rather than genuine protein signal. Bars indicate average values, error bars their SD, values above each bar indicate summed intensities (n = 2 technical replicates). (**B**) Number of quantified proteins across the indicated methods (see table below for mode, gradient length, RI region and input). DDA used an isolation window of 0.5 *m/z* with a MS2 IT of 8 ms, while DIA methods used an isolation window of 8 *m/z* with a MS2 IT of 120 ms. DIA120 uses a 120 min active gradient and HM-DIA does not include the RI region. The number of quantified proteins is given for TMT RI quantification before/after filtering (co-isolation purity ≥ 70%, TMT RI signal-to-noise ratio ≥ 20 and TMT RI resolution ≥ 45,000) and for TMTc quantification (n = 2 technical replicates). (**C**) Box plots showing the normalized (weighted average) log2 fold change (FC) for human (blue) and yeast (orange) proteins in channels 131N or 135N based on TMTc quantification across the indicated DIA and DDA methods. The horizontal center line represents the median, the lower and upper hinges correspond to the first and third quartiles (25^th^ and 75^th^ percentiles, defining the interquartile range (IQR)) and the whiskers extend to the furthest data points within 1.5x IQR from the hinges. Outliers are omitted from the visualization for clarity. Values above each box represent the corresponding median. Horizontal dashed lines indicate theoretical/expected log2 FC: 0 for both human and yeast in channel 131N, and 0 for human and 2.32 for yeast in channel 135N (n = 2 technical replicates).

We next compared the number of quantified proteins across the above-mentioned methods at the specified inputs. TMT RI quantification was evaluated both before and after stringent filtering (co-isolation purity ≥ 70%, TMT RI S/N ≥ 20, TMT RI resolution ≥ 45,000), alongside TMTc-based quantification (Figure 5B). Across all methods, TMTc quantification yielded substantially fewer quantified proteins than unfiltered TMT RI quantification. Compared to filtered TMT RI data, however, TMTc achieved comparable depth for the tested DIA methods, while DDA clearly favored TMT RI quantification even after quality filtering (e.g., 5485 vs. 4998 vs. 556 quantified proteins for 50 ng input). This reduction in TMTc coverage likely reflects, at least in part, the stricter processing criteria applied to TMTc, which require detectable complementary ion signal across 131N and 135N per peptide-spectrum match (PSM). We additionally note that the DIA methods used here employ a broader isolation window, longer ITs, and a wider MS1 scan range than the two-proteome TMTpro 18-plex experiment used for our co-isolation interference benchmarking (Figure 3). This increases co-isolation and, after quality filtering, drastically reduces the number of quantifiable proteins. We therefore present this comparison primarily to show the trade-off between interference resistance and proteome coverage, rather than a controlled head-to-head benchmark.

Finally, we assessed quantitative accuracy by comparing the normalized (weighted average) log2 FC for human and yeast proteins in the 131N and 135N channels across all DDA and DIA methods (Figure 5C). Measured ratios closely matched the expected values across all methods for both human and yeast proteins. This confirms that TMTc-based quantification achieves high accuracy even under co-isolation conditions. Yeast proteins showed a notably broader spread in the 131N channel compared to 135N across methods. We attribute this primarily to the lower absolute signal intensity in channel 131N. At lower ion abundance, the complementary ion signal approaches the detection and noise-threshold limits more often, increasing the relative contribution of noise to the measured ratio.

### Quantitative accuracy across LFQ, TMT RI and TMTc quantification strategies

Having benchmarked proteome coverage and interference resistance across the TMT RI and TMTc approaches (Figure 5), we eventually assessed quantitative accuracy across all tested quantification strategies used in this study: DIA-LFQ, DDA-TMT RI, DIA-TMT RI, and DIA-TMTc, using matched human-yeast mixtures at defined ratios (Figure 6).

**Figure 6:**
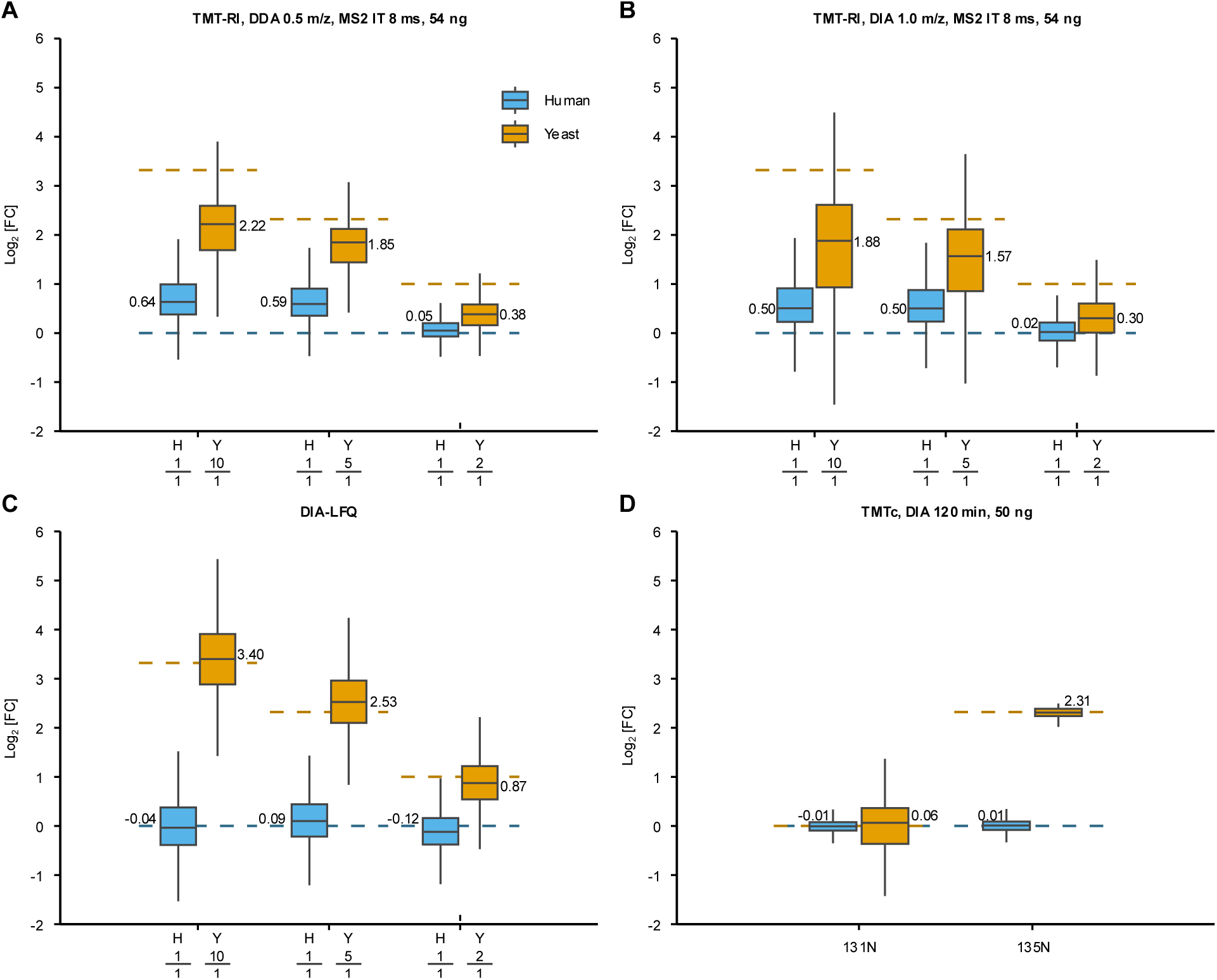
Comparison of quantification accuracy across sample designs and quantification methods (LFQ, TMT RI and TMTc). Box plots showing the log2 fold change (FC) for human (blue) and yeast (orange) proteins between the indicated sample mixtures. (**A, B**) Log2 FC for TMT reporter ion (RI)-based quantification of the pooled sample using a DDA (0.5 *m/z*, MS2 IT 8 ms, 54 ng; A) and DIA (1 *m/z*, MS2 IT 8 ms, 54 ng; B) method for the shown comparisons (filtered for co-isolation purity ≥ 70%, TMT RI signal-to-noise ratio ≥ 20 and TMT RI resolution ≥ 45,000; n = 3 technical replicates). (**C**) As in (A, B), but for label free quantification (LFQ) in DIA (n = 3). (**D**) Log2 FC (weighted average normalization) for human (blue) and yeast (orange) proteins in TMT channels 131N and 135N, quantified by TMTc using a DIA method (8 *m/*z, MS2 IT 120 ms, 120 min gradient, 50 ng; n = 2 technical replicates). The horizontal center line represents the median, the lower and upper hinges correspond to the first and third quartiles (25th and 75th percentiles, defining the interquartile range (IQR)) and the whiskers extend to the furthest data points within 1.5x IQR from the hinges. Outliers are omitted from the visualization for clarity. Values corresponding to each box represent the median. Horizontal dashed lines indicate theoretical/expected log2 FC.

For TMT RI quantification, we compared DDA (Figure 6A) and DIA (Figure 6B) methods at 54 ng input, evaluating three sample mixtures with expected log2 FC of 3.32, 2.32 and 1 for yeast (human log2 FC = 0 throughout). Both methods recovered the expected direction of yeast FCs, but measured ratios were compressed relative to the theoretical values in all three comparisons (e.g., median log2 FC of 2.22 vs. expected 3.32 for the log2 FC of 3.32 in DDA and 1.88 vs. 3.32 in DIA). For human proteins, measured ratios closely matched the expected log2 FC of 0 in the mixture with the lowest yeast spike-in (median log2 FC of 0.05 for DDA and 0.02 for DIA) but increasingly deviated above 0 as the amount of spiked-in yeast in these specific channels increased (median log2 FC of 0.64 for DDA and 0.50 for DIA at the highest yeast spike-in). Both effects, compression of the yeast ratio and inflation of the nominally flat human ratio, are consistent with co-isolation of yeast and human precursors within the same isolation window: as more yeast peptide is present, a larger fraction of the fragmented ion population originates from yeast, which pulls the quantified human ratio upward toward the dominant yeast signal while simultaneously pulling the yeast ratio itself downward toward the constant human background, as previously discussed and shown in Supplementary Figure 3. DIA showed somewhat less-inflated human ratios than DDA but simultaneously suffered from higher ratio compression for the yeast ratios.

DIA-LFQ quantification (Figure 6C) of the TMT-labeled samples (measured as single channels prior to pooling) showed no systematic ratio compression, with human and yeast ratios close to their expected values across all three comparisons (e.g., yeast median log2 FC of 3.40 vs. expected 3.32 in the 1:10 comparison; human medians of −0.04, 0.09 and −0.12 across the three comparisons, all close to the expected 0). This improved accuracy in comparison to TMT-RI quantifications reflects the absence of co-isolation interference in LFQ, since each channel is acquired and quantified individually via DIA before TMT pooling, rather than as part of single multiplexed precursors alongside the other channels. This is in agreement with He *et al*.^19^, who similarly reported higher quantitative accuracy for LFQ compared to TMT RI-based quantification. However, this strategy comes at the cost of lower throughput.

TMTc-based quantification (Figure 6D) also showed close agreement with expected values for both channels and species: median log2 FC 0.01 and −0.01 for human proteins in channels 135N and 131N, respectively (expected 0), and 2.31 and 0.06 for yeast in 135N and 131N (expected 2.32 and 0, respectively). This confirms that TMTc retains high accuracy for complex, co-isolated samples, consistent with its resistance to co-isolation interferences, as discussed for Figure 5. As noted previously, this accuracy is achieved at the cost of reduced proteome depth and precision suffers specifically in the low intensity 131N channel for yeast proteins.

Taken together, these results illustrate a consistent trade-off across the three quantification strategies: LFQ offers interference free quantification but at a reduced throughput, since each sample must be measured in a separate run. TMT RI quantification enables high multiplexing and throughput but is susceptible to co-isolation-driven ratio distortion – manifesting as both compression and inflation – regardless of whether data are acquired by DDA or DIA. TMTc combines the multiplexing advantage of isobaric labeling with near-LFQ-level accuracy by exploiting interference-resistant complementary ions, but only at strongly reduced plexing capacity and proteome depth. Together, these findings position TMTc as a complementary rather than a replacement for RI-based TMT: suited for applications prioritizing quantitative accuracy for a small number of conditions, while conventional TMT RI quantification remains preferable when broad proteome coverage and high sample multiplexing are the primary goals.

## Conclusion

In this study, we systematically evaluated whether the complementary strengths of DIA and TMTpro multiplexing can be combined to improve the scalability of low-input proteomics, such as in SCP. DIA is attractive because its systematic precursor sampling can increase identification depth and data completeness relative to the stochastic precursor selection of DDA, while TMT provides the throughput required to analyze large numbers of samples or single cells. Our results, however, show that transferring these advantages into accurate RI quantification is constrained by precursor co-isolation resulting in ratio compression and inflation under the tested low-input but highly complex sample conditions. Although DIA-TMT initially produced more protein identifications than DDA-TMT, its fixed isolation windows captured interfering precursors more frequently, particularly at wider windows and higher sample loads. Narrower windows improved precursor co-isolation purity but increased cycle time, reduced ion statistics and diminished identification depth. Thus, the central advantage anticipated for DIA – broader and more reproducible sampling – was offset by the acquisition and filtering requirements needed for reliable TMT quantification.

After stringent filtering for co-isolation purity, RI S/N and resolution, DDA and DIA achieved broadly comparable quantitative performance, although neither approach fully eliminated ratio distortion in the two-proteome interference model. Both acquisition modes recovered the direction of regulation, but underestimated true yeast FCs and, at high spike-in levels, inflated ratios of the nominally constant human background. These effects were substantially reduced in the single-proteome benchmark, demonstrating that the principal limitation is co-isolation in complex mixtures rather than acquisition mode alone. Importantly, filtering removed a large fraction of DIA identifications and thereby largely erased its initial gain in coverage. Consequently, DIA-TMT cannot be considered intrinsically superior to DDA-TMT under the tested conditions: once quantitative reliability is prioritized, both methods can perform similarly, but both remain imperfect, and DIA no longer retains a clear advantage in proteome depth.

Because quantification on MS3 level is not possible on the Orbitrap Astral mass spectrometer, alternative strategies that decouple quantification from precursor co-isolation – such as TMTc – are of particular interest. TMTc quantification reproduced the expected ratios under co-isolation conditions and approached the accuracy achieved by DIA-LFQ. This gain, however, came with substantial practical limitations. Our implementation was restricted to three well-separated TMTpro channels since deconvolution of isotopic envelopes and MS resolving power limit robust discrimination of the more closely spaced complementary-ion masses required for higher plexing. TMTc also yielded fewer quantified proteins than RI-based workflows, with a particularly pronounced loss for DDA, and precision deteriorated for low-intensity complementary-ion signals. TMTc therefore represents an accuracy-oriented solution for a limited number of conditions rather than a direct replacement for high-plex RI quantification. DIA-LFQ provided the most interference-resistant quantification and recovered the expected ratios without systematic compression, but each sample required an individual LC-MS/MS run. Its high quantitative fidelity and proteome depth are therefore balanced by lower throughput and greater instrument-time requirements as cohort size increases.

Future progress will require methods that preserve the high multiplexing capacity of TMT or non-isobaric tag alternatives^28,29^ while reducing interference. Key developments include faster and more sensitive instrumentation, higher effective resolving power for complementary ions, acquisition schemes that adapt isolation windows to precursor density and purity, improved ion-statistical efficiency at low input, and computational approaches that identify or correct interference without relying on highly restrictive post-acquisition filters.

In conclusion, we believe that with current state-of-the-art instrumentation, TMT and LFQ are likely to occupy complementary positions in single-cell proteomics. TMT RI approaches will remain advantageous when sample throughput, cohort size and cost per sample/cell are dominant considerations, particularly for screening designs in which robust detection of the presence and direction of regulation is more important than recovering the exact magnitude of every FC. LFQ will be preferred when maximal proteome depth, quantitative accuracy, dynamic-range fidelity and avoidance of ratio compression are essential, especially for mechanistic studies, subtle biological effects or smaller cohorts in which instrument time is less limiting. TMTc may bridge these regimes for experiments with few conditions that demand multiplexing and high accuracy, provided that reduced depth and plexing are acceptable. Overall, our data establish DIA-TMT as technically feasible on the Orbitrap Astral mass spectrometer platform but also define the boundary conditions for its use: DIA and DDA can support comparable TMT quantification after rigorous quality control, whereas the optimal choice among reporter-ion TMT, TMTc and LFQ should be determined by the required balance of throughput, coverage and quantitative accuracy.

## Methods

### Commercial protein digest standards

HeLa (88328; Thermo Fisher Scientific) and yeast (V7461, digest; Promega) protein digests standards were used. The peptide standards were dissolved in 100 mM tetraethylammonium bromide (TEAB) (102699716, Sigma-Aldrich) to a final concentration of 1 µg/µL or 2 µg/µL each. Subsequently, yeast and human peptides were combined at pre-defined ratios specified in the respective result section.

### K562 cell culture and in-solution digest

To achieve higher inputs for labeling, we additionally cultured K562 cells in RPMI 1640 medium for bulk digests. The medium was supplemented with 10% fetal bovine serum (10270; Thermo Fisher Scientific), 100x L-glutamine (250030-024; Thermo Fisher Scientific) and 1x streptomycin-penicillin (P0781-100ML; Sigma-Aldrich). Cells were cultured at 37 °C, 5% CO2 and harvested at 80% confluence by centrifugation for 3 min at 200x g and 4 °C, followed by washing with PBS. Cells were pooled to 1x 10E7 cells, snap frozen and stored at −70 °C until further use. Snap-frozen K562 cells were lysed in a lysis buffer consisting of 10 M urea in Tris-HCl buffer using sonication. Subsequently, the lysed K562 cells were reduced with 10 mM dithiothreitol (DTT) and alkylated with 20 mM iodoacetamide; the alkylation reaction was quenched by adding DTT to a final volume corresponding to 50% of that used during reduction. Both samples were diluted to 6 M urea with 100 mM Tris-HCl before digestion with Lys-C overnight on a thermoshaker at 37 °C, followed by trypsin digestion for 2 h on a thermoshaker at 37 °C. Samples were desalted on a Sep-Pak C18 cartridge (WAT054955; Waters), vacuum-dried and resuspended in TEAB.

### Tandem mass tag labeling procedure

TMTpro 18-plex (A52045; Thermo Fisher Scientific) was dissolved in anhydrous acetonitrile according to the manufacturer’s instructions (c = 25 µg/µL) and stored at −70 °C until further use. Labeling, quenching, pooling and clean-up were performed identically for all sample sets described below unless noted otherwise: the respective label reagent was added to each sample with a w:w peptide-to-TMT reagent ratio ranging from 1:5-1:10 and the labeling reaction was conducted for 1 h at room temperature. A small aliquot of each sample was subsequently taken to assess labeling efficiency, and the remaining samples were kept at −70 °C until further processing. Samples were then quenched with methylamine to a final concentration of 0.4 M, following an established protocol^30^. Samples were vacuum-dried and reconstituted in trifluoroacetic acid (28903; Thermo Fisher Scientific) prior to pooling at equal volumes. The pooled sample was cleaned up using the C18 StageTips (98-0604-0220-7EA; CDS Analytical)^31,32^, vacuum-dried again and reconstituted in 0.1% trifluoroacetic acid, 0.015% n-dodecyl β-D-maltoside. A diluted sample at a concentration of 50 ng/µL was prepared for multiple injections into the LC-MS/MS system.

### Two-proteome mix for TMT RI quantification

Human:yeast mixtures from commercial protein digests were prepared according to the schema in Figure 1B, where channels 126, 127N, 127C, 129C, 131C and 133C contained no peptides, but only 100 mM TEAB. The experimental design resulted in five groups for the TMTpro 18 set: H0:Y0, H1:Y0, H1:Y0.1, H1:Y0.2 and H1:Y1 with six intra-sample replicates for the zero controls and three intra-sample replicates for each of the remaining conditions. Prior to pooling all channels, we measured an aliquot of each sample in DIA-LFQ to compare the quantitative performance to TMT RI.

### Two-proteome mix for TMTc quantification

For the TMTc approach, the standard HeLa and yeast digest standards described above were labeled using only three channels of the TMTpro 18-plex set: 126, 131N and 135N (supplementary Figure 1B). Peptides were combined at H0:Y0, H1:Y0.05 and H1:Y0.25 respectively. The TMT labeling procedure was otherwise identical to that described above.

### Human-only sample

For higher inputs, the in-house cultured and digested K562 peptides were labeled analogously using TMTpro 18-plex reagent, following the labeling, quenching, pooling and clean-up procedure described above. Channels were assigned to four groups: H0, H1, H5 and H10 (supplementary Figure 1A). This resulted in six intra-sample replicates for the zero controls and in four intra-sample replicates for each of the remaining conditions.

### LC-MS/MS analysis

Samples were separated on the Vanquish™ Neo UHPLC (VN-S10-A01; Thermo Fisher Scientific) at 50 °C in trap-and-elute mode, using an Aurora® Ultimate™ C18 UHPLC column (1.7 µm, 25 cm x 75 µm) (AUR5-25075C18-XT; IonOpticks), with an integrated emitter. Peptides were eluted with a 60 min active gradient from 2% to 40% mobile phase B (80% acetonitrile in 0.08% FA) at a flow rate of 300 nL/min. For the DIA120 method (Figure 5) we doubled the active gradient to 120 min. The LC was coupled to the Orbitrap Astral mass spectrometer for MS measurement, equipped with a FAIMS Pro Duo interface and an EASY-Spray source (all Thermo Fisher Scientific). The FAIMS interface was operated at a compensation voltage of −48 V for both DDA and DIA experiments. An electrospray voltage of 1900 V was applied to the emitter, and the ion transfer tube was operated at 280 °C.

Measurement of the single channels from the two-proteome TMT-mix prior to pooling was conducted in DIA mode: MS1 scans were acquired at a resolution of 240,000 in a range of 400-900 *m/z*. The normalized automatic gain control target (AGC) was set to 500% with an IT of 100 ms. Precursors were isolated in an isolation window of 5 *m/z*, fragmented by HCD at a normalized collision energy of 29% before MS/MS. Fragment ions were collected over the scan range of 110-2000 *m/z* for an IT of 10 ms or until reaching the AGC target of 800%.

For the two-proteome TMT RI mix, DDA-TMT experiment parameters were adapted from one of the first studies applying TMT on the Orbitrap Astral mass spectrometer by Keele *et al.*^33^. The DDA parameters were as follows: MS1 scans were acquired over a range of 350-1350 *m/z* at an Orbitrap resolution of 60,000. The AGC was set to 200% with a maximum IT of 100 ms. Only precursors with a purity of greater than 70% and a charge state between 2 and 6 were selected for fragmentation in MS2. Precursors were isolated for fragmentation using a 0.5 *m/z* window, with a maximum IT of either 8 or 20 ms and AGC was set to standard. Fragmentation of precursors was conducted by HCD at a NCE of 35% before MS/MS measurement in the Astral mass analyzer.

For the two-proteome TMT RI mix, DIA-TMT experiments were recorded on the Orbitrap with a resolution of 240,000 from 400-800 m/z for MS1. Maximum IT and AGC were identical as in the DDA experiments. Precursors were isolated in isolation windows of 0.5, 1 or 2 *m/z*, fragmented by HCD at a NCE of 25% and fragment ions were collected over a scan range of 110-2000 *m/z* for a maximum IT of 8, 20 or 40 ms or until reaching the AGC target of 800%. Loop control was enabled and set to 0.6 s. For both DIA and DDA experiments the TMT mode was activated.

If not mentioned differently, DIA and DDA parameters for the measurement of the human-only TMT sample as well as DDA-TMTc samples were identical to those described above for the two-proteome TMT RI mix.

For DIA-TMTc experiments, MS1 spectra were recorded on the Orbitrap at a resolution of 120,000 from 350-1350 *m/z*, with a maximum IT of 100 ms and an AGC target of 300%. Precursors were isolated in windows of 8 *m/z* with an overlap of 1 *m/z* and fragmented by HCD at a NCE of 29%. Fragment ions were collected for a maximum IT of 120 ms or until reaching an AGC target of 2000%, and MS2 spectra were recorded from 120-2550 *m/z*. Loop control was enabled and set to 0.6 s. TMT mode was again activated for both DDA and DIA experiments. Additionally, we tested a narrower MS2 scan range of 255-2550 *m/z*, which excluded the TMT RI region.

### Data analysis

#### Raw file processing

Unless stated otherwise, the RAW data were analyzed with the CHIMERYS^34^ node using the prediction model inferys (v4.7.0) in Proteome Discoverer (v.3.2, Thermo Fisher Scientific). The RAW files were processed individually. Static modifications were defined as TMTpro 18 tags on lysine residues and peptide N-termini (+304.207 Da), and carbamidomethylation on cysteine residues (+57.021 Da). Oxidation of methionine (+15.995 Da) was set as variable modification. The number of missed cleavages permitted was 2. Fragment mass tolerance was kept default at 20 ppm. False discovery rate (FDR) was set to 1% for PSM for high confidence. The same FDR was applied at the peptide and protein level. Only unique peptides were used for protein quantification. To assess the labeling efficiency (LE), the same CHIMERYS settings were applied, except that TMTpro 18 tags were set as variable modifications on lysine and N-terminus.

Furthermore, selected DIA-files were analyzed with DIA-NN (v2.2.0 Academia) and Spectronaut (v20.1.250624.92449) with the same parameters as in the CHIMERYS search, if not stated otherwise. For the DIA-NN searches, automatic inference of precursor and fragment mass tolerance was enabled. Match between runs was disabled and precursor FDR was set to 1%. In the Spectronaut searches, mass tolerances were determined dynamically. Method evaluation was enabled and each file was defined as separate condition to avoid cross-file matching. For both Spectronaut and DIA-NN, TMTpro tags were set as fixed modification on N-termini and lysine residues.

DIA data of the single channel injections was analyzed with Spectronaut (v20.1.250624.92449) for LFQ, with factory settings if not stated otherwise. TMTpro tags were set as fixed modification on N-termini and lysine residues.

For TMTc quantification, corresponding RAW files were first searched in FragPipe^35,36^ (v.23.0) with MSFragger (v.4.3) with TMT-18 as fixed modification on lysine and peptide N-termini. Methionine oxidation and protein N-terminal acetylation were included as variable modifications. DIA files were searched in library-free mode and results were filtered at a 1% FDR on the peptide level. For DDA files a 1% FDR cutoff was applied at the PSM level.

All above described searches were performed against the human proteome (Uniprot: UP000005640, reviewed, 20,650 protein entries, downloaded 02.12.2024), the yeast proteome (Uniprot: UP000002311, reviewed, 6,733 protein entries, downloaded 05.02.2025) and the CRAPome^37^ (118 entries).

### Post-processing and annotation of TMT RI data using an in-house Python script

We developed a python-based tool called READ – short for **R**eporter Ion **E**xtractor and **A**nnotation **D**irector – to orchestrate quantification, purity calculation, and additional quality control. READ supports mass spectra in both Thermo RAW or mzML format^38^ as input and additionally requires an identification file with PSMs or precursors from either CHIMERYS, Spectronaut, or DIA-NN. Thermo RAW files are automatically converted to mzML format using ThermoRawFileParser v1.4.5^39^ and then read with pyteomics^40,41^. TMT RI are subsequently quantified with OpenMS using the IsobaricAnalyzer tool^42^. Precursors and PSMs are matched to their respective quantified RI via their corresponding MS2 spectra which for CHIMERYS is carried out via the reported MS2 scan number. Since Spectronaut and DIA-NN do not report a scan number, the MS2 scan is identified by searching for the scan that is closest to the identified precursor in terms of *m/z* and retention time.

Then the parent MS1 scan is identified to calculate precursor co-isolation purity. Purity is calculated as follows:

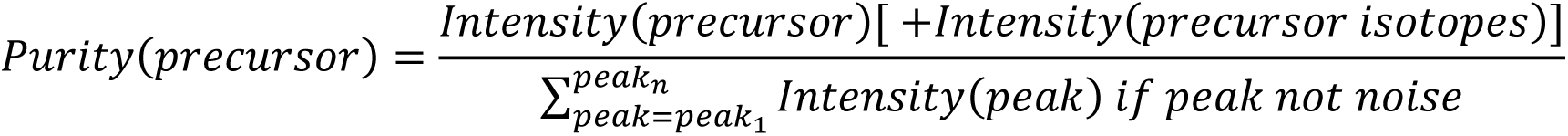

The purity is calculated as the intensity of the precursor (plus optionally the intensity of all precursor isotopes) divided by the sum of all the intensities of all peaks *peak1* to *peakn* in the isolation window. The intensity of peaks that are classified as noise is not summed up. We classify peaks as noise if they are below *N*% of the precursor intensity where *N* is a user-definable parameter and in our case 10% (default).

For further quality control we calculate S/N per condition, RI resolution, and optionally subtract reporter noise from reporter signal. All of this is based on the output of the Resolution GUI tool developed by Keele and colleagues^33^ which can be passed to the script as an additional input. Please note that we used a slightly adapted version of the tool that supports DIA data.

READ is publicly available under a permissive open-source license via the GitHub repository https://github.com/hgb-bin-proteomics/READ with detailed documentation available at https://hgb-bin-proteomics.github.io/READ. READ runs on any operating system, and we also offer compiled binaries for Windows users that come with a graphical user interface.

### TMTc data post-processing

Resulting PSM tables from the FragPipe search were subsequently processed with a published Python script^25^ for TMTc quantification. The Python script was adapted to our experimental setup as follows: The complement ion of channel 126 was excluded from quality filtering, as this channel was intentionally left empty. The intensity threshold for complementary ions of 131N and 135N was lowered to 2,500 and for the M+1 peak to 250, due to the lower input in our approach. PSM ranking at the peptide level was only based on 131N and 135N. All other filtering and aggregation logic followed the original published implementation in the Python scripts.

### Downstream data analysis

The labeling efficiency for each channel was calculated as the ratio of observed TMT modifications on the corresponding PSMs to the total number of possible TMT modification sites. This calculation was performed for the total number of PSMs and additionally assessed separately for yeast and human PSMs.

For downstream analysis the annotated PSM and proteins tables were processed in R (v4.5.0) within RStudio (v2025.05.0+496) using base R or the packages tidyverse^43^, cowplot^44^, patchwork^45^ and ggplot2^46^.

Downstream processing of the TMT RI data included removing contaminants and replacing zeros with NAs. Furthermore, the CV and ratios between conditions were only calculated when a protein was quantified in all intra-sample replicates per technical replicate.

For downstream analysis of the TMTc data, contaminants were removed, and noise subtraction was performed by calculating, separately for human and yeast proteins, the median intensity plus one standard deviation in the empty 126 channel. The corresponding species-specific value was then subtracted from the 126, 131N and 135N intensities of that species. Abundance ratios were calculated by dividing each corrected channel intensity by a species-specific weighted average, defined as the sum of the two corrected channel intensities (131N and 135N) divided by the sum of their expected relative abundances (i.e., the nominal channel mixing ratios: 1:1 for human and 1:5 for yeast, reflecting the H1:Y0.05/H1:Y0.25 sample design). Ratios were log2-transformed for all subsequent analyses.

DIA-LFQ protein data was imported into R and 0 values were converted to NA. For each sample group, the mean log2 intensity and CV across replicate channels were calculated only when a protein was quantified in all replicates of that group. Pairwise log2 FC between groups were calculated as the difference of the respective group means.

If not stated otherwise, visualizations were generated in R using ggplot2.

## Data availability Code availability

READ – the python-based tool for quantification, purity calculation, and additional quality control – is available via the GitHub repository https://github.com/hgb-bin-proteomics/READ.

The adapted version of the Resolution GUI tool originally developed by Keele *et al.*^33^ that we used for our analyses is available at https://github.com/hgb-bin-proteomics/TMT_Resolution_GUI.

## Supplemental data

This article contains supplementary data.

## Acknowledgements

All LC-MS/MS analyses were performed on the Vienna BioCenter Core Facilities instrument pool. We thank Tabiwang Arrey, Bernard Delanghe and Dominic Hoch from Thermo Fisher Scientific for fruitful discussions and their constructive feedback. We would like to thank Isabella Burger for proofreading the manuscript.

We thank Dina L. Bai and Tian Zhang for their original implementation of the Resolution GUI tool, for their help in using the tool, and their support in developing and sharing the adapted version of the tool that was used in this manuscript. We furthermore thank Dasha Pototska for designing the READ logo.

Troubleshooting and finetuning of custom R scripts was supported by Gen-AI using Claude Sonnet 5 in Abacus (abacus.ai). While preparing this work, Abacus was also used to assist in editing and improving the clarity and language of the text. The authors carefully reviewed and edited all AI-assisted content, taking full responsibility for the publication’s content.

## Funding and additional information

This work was supported by the infrastructure funding 4th call 2022/01 (AT-SCP) of the Austrian Research Promotion Agency (FFG) as well as by the FWF projects 10.55776/PAT4800425, 10.55776/PAT4142423 and 10.55776/PAT2059025. For the purpose of open access, the authors have applied a CC BY public copyright license to any Author Accepted Manuscript version arising from this submission.

## Author contributions

T.T. and M.M. conceptualized the study. T.T. performed experiments, analyzed the data and wrote the manuscript. M.M. and K.M. supervised the study. M.J.B wrote and implemented custom Python scripts (READ) for downstream data processing. T.T., M.J.B., V.D., K.M. and M.M. revised and agreed on the manuscript.

