## Supplemental Figures for "Making the Impossible Possible: Uniting TMT and DIA for Scalable Low-Input-Proteomics"

### Content

**Supplementary Figure 1:** Experimental designs for the single-proteome benchmark and the TMTc benchmark.

**Supplementary Figure 2:** Effect of filtering stringency on protein identification, precision and quantitative accuracy.

**Supplementary Figure 3:** Ratio compression and inflation scale with yeast spike-in level.

**Supplementary Figure 4:** Search software choice has minor influence on TMT RI-based quantification.

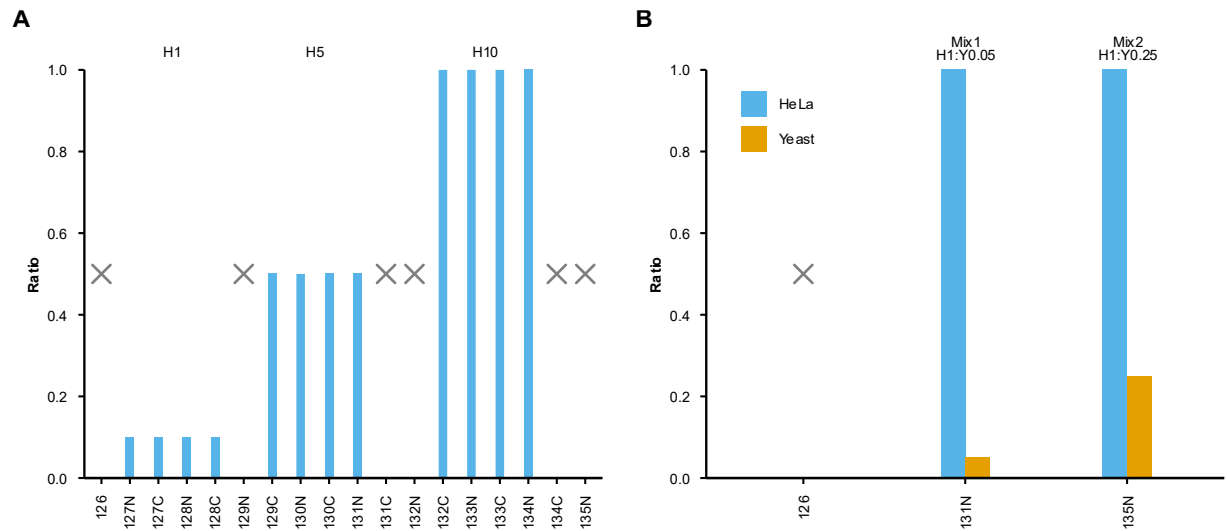

**Supplementary Figure 1: Experimental designs for the single-proteome benchmark and the TMTc benchmark.** (A) Relative loading scheme for the single-proteome human benchmark comprising four groups across the TMTpro 18-plex: H0 (zero control, six intra-sample replicates, marked "x"), H1, H5 and H10 (relative loading ratios of 1×, 5× and 10×, respectively; four intra-sample replicates each). (B) Labeling scheme for the TMTc benchmark, using three channels of the TMTpro 18-plex: channel 126 (H0:Y0, empty control), channel 131N (Mix1, H1:Y0.05) and channel 135N (Mix2, H1:Y0.25), combining a constant HeLa (human) background with increasing yeast spike-in levels.

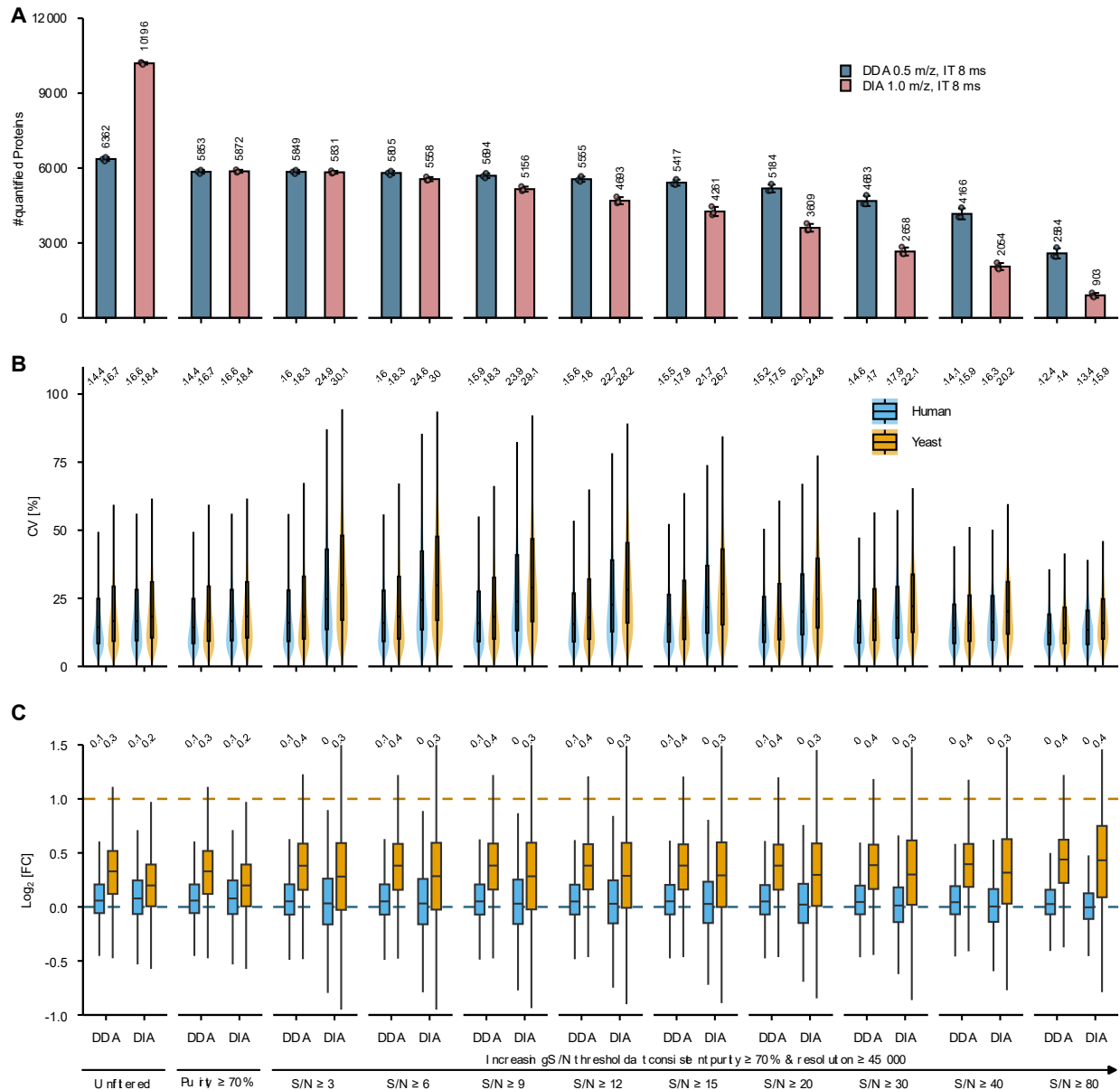

**Supplementary Figure 2: Effect of filtering stringency on protein identification, precision and quantitative accuracy.** Representative DDA (0.5 m/z, MS2 IT 8 ms, 54 ng) and DIA (1 m/z, MS2 IT 8 ms, 54 ng) methods were evaluated across increasing filter stringency: unfiltered, co-isolation purity ≥ 70% alone, and purity ≥ 70% combined with TMT reporter ion (RI) resolution ≥ 45,000 and increasing TMT RI S/N thresholds (≥3, 6, 9, 12, 15, 20, 30, 40, 80) (n = 3 technical replicates). **(A)** Number of quantified proteins at each filter step. **(B)** Violin plots of coefficient of variation (CV [%]) for human (blue) and yeast (orange) proteins at each filter step; numbers above each violin indicate median CV. **(C)** Box plots of log<sub>2</sub> fold change (FC) for human (blue) and yeast (orange) proteins between Mix2 and Mix1 at each filter step; values above each box pair indicate medians. Horizontal dashed lines indicate theoretical/expected log<sub>2</sub> FC (0 for human, 1 for yeast).

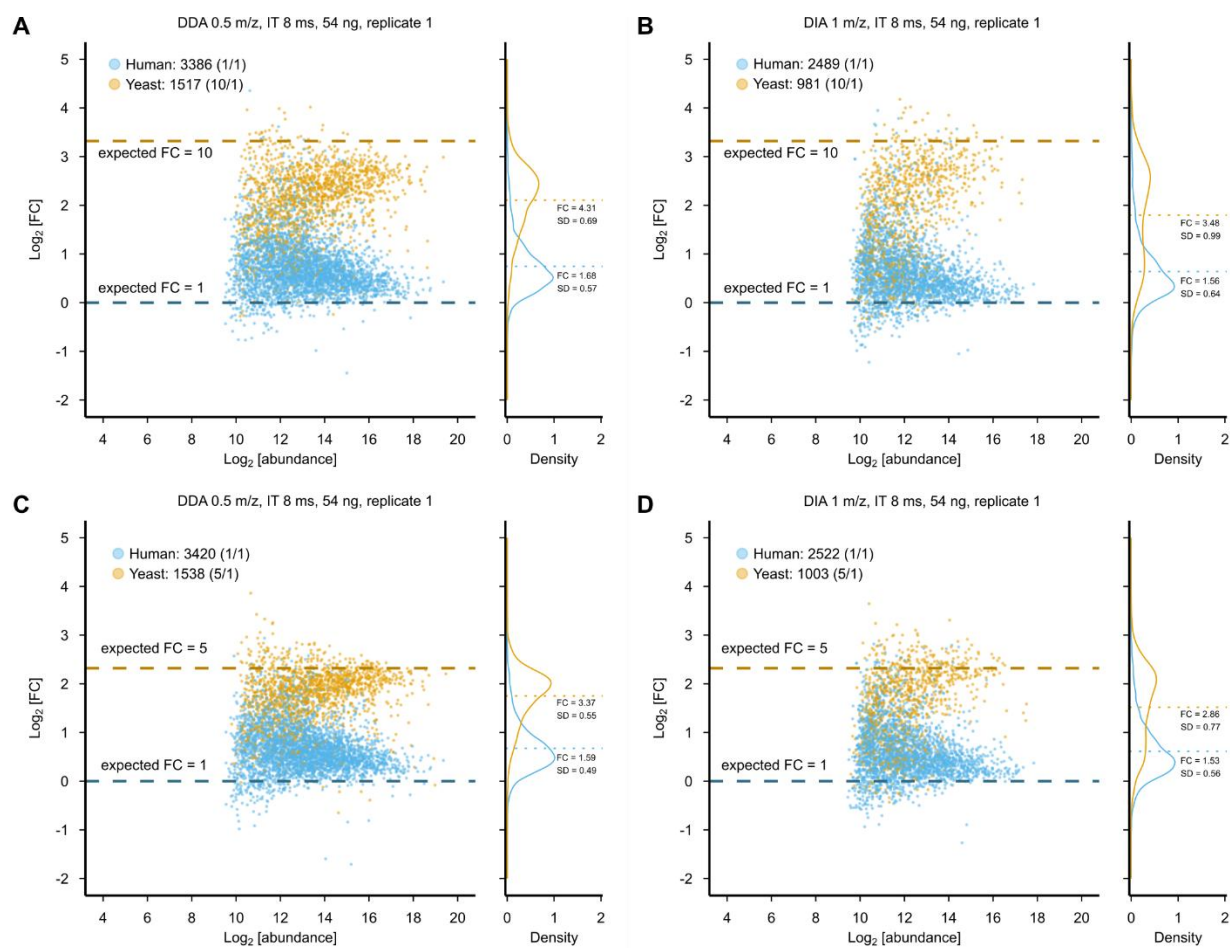

**Supplementary Figure 3: Ratio compression and inflation scale with yeast spike-in level.** Representative Bland-Altman plots from one technical replicate of a DDA (0.5 *m/z*, MS2 IT 8 ms, 54 ng; **A**, **C**) and DIA (1 *m/z*, MS2 IT 8 ms, 54 ng; **B**, **D**) method, showing measured $\log_2$ fold changes (FC) for human (blue) and yeast (orange) proteins between Mix3/Mix1 (expected yeast $\log_2$ FC = 3.32; **A**, **B**) and Mix3/Mix2 (expected yeast $\log_2$ FC = 2.32; **C**, **D**), where the theoretical $\log_2$ FC is 0 for human (dashed blue line) and as indicated for yeast (dashed orange line). Legend indicates the number of quantified proteins for each species and the corresponding expected linear fold change (in parentheses). Density plots on the right show the distribution of measured $\log_2$ FC, with the median FC and standard deviation (SD) indicated for each species ($n = 3$ intra-sample replicates).

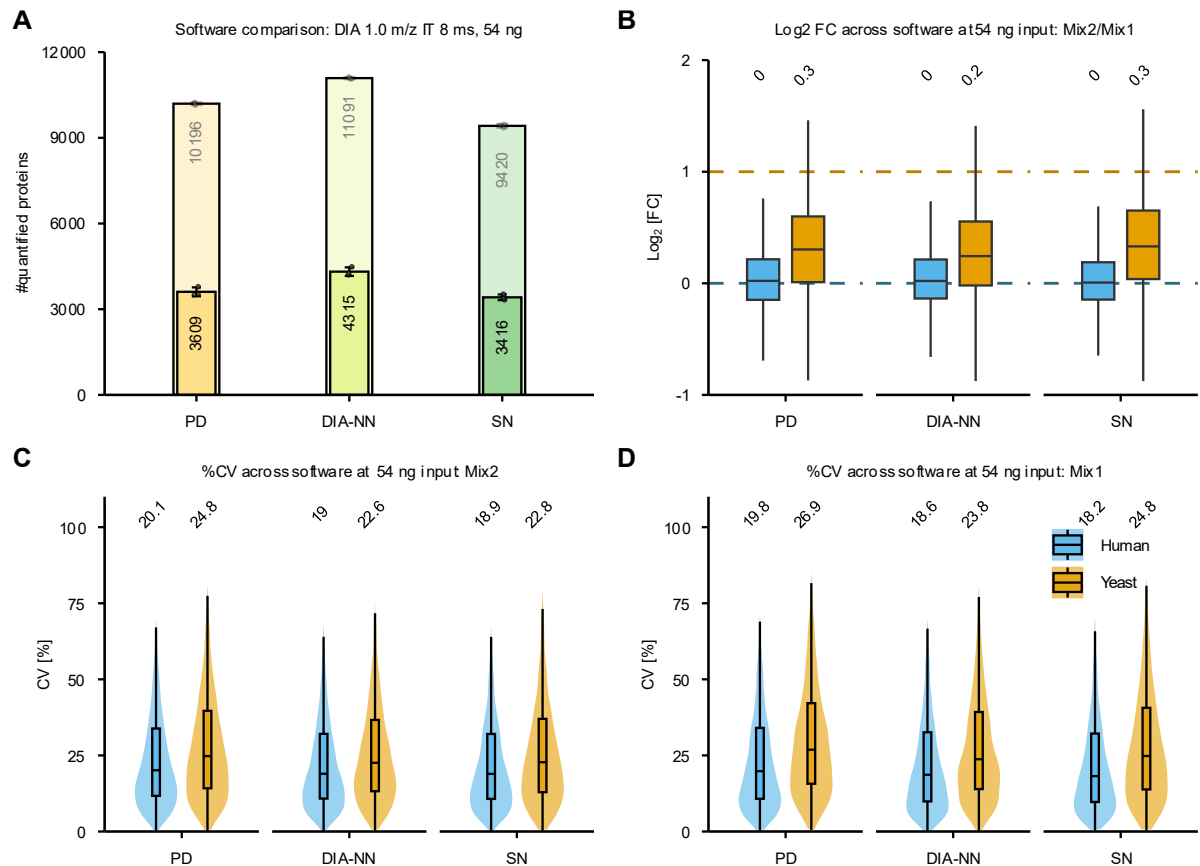

**Supplementary Figure 4: Search software choice has minor influence on TMT RI-based quantification.** (A) Number of quantified proteins before (light, top values) and after (solid, bottom values) filtering (co-isolation purity $\geq 70\%$, TMT reporter ion (RI) signal-to-noise ratio $\geq 20$ and TMT RI resolution $\geq 45,000$) for a representative DIA method (1 $m/z$ isolation window, MS2 IT 8 ms, 54 ng) searched with Chimerys in Proteome Discoverer (PD), DIA-NN and Spectronaut (SN); bars indicate average values, error bars their standard deviation, $n = 3$ technical replicates. (B) Box plots showing the log<sub>2</sub> fold change (FC) for human (blue) and yeast (orange) proteins between Mix2 and Mix1 across the three search software packages. The horizontal center line represents the median, the lower and upper hinges correspond to the first and third quartiles (25<sup>th</sup> and 75<sup>th</sup> percentiles, defining the interquartile range (IQR)) and the whiskers extend to the furthest data points within 1.5x IQR from the hinges. Outliers are omitted from the visualization for clarity. Values above each box represent the corresponding median. Horizontal dashed lines indicate theoretical/expected log<sub>2</sub> FC (0 for human and 1 for yeast). (C, D) Violin plots show the distribution of coefficients of variation (CV [%]) for human (blue) and yeast (orange) proteins for Mix2 (C) and Mix1 (D) across the three search software packages. Violin width reflects the relative density of data points at a given CV value. Overlaid boxplots indicate the median (center line) and IQR (25<sup>th</sup>-75<sup>th</sup> percentiles, box limits); outliers are omitted for clarity. Numbers above each violin show the respective median CV ($n = 3$ technical replicates).
